# Genomic signature of repeated transitions to diurnality in spiders

**DOI:** 10.1101/2025.11.23.690066

**Authors:** Chao Tong, Zheng Fan, Luyu Wang, Zhisheng Zhang

## Abstract

Repeated transitions to diurnality represent a major behavioral shift in spiders, yet their genomic underpinnings remain largely unknown. Here, we assembled high-quality genomes for nine diurnal spiders using long-read sequencing and compiled diel activity phenotypic data through a combination of meta-analysis and field assessments. By integrating all publicly available spider genomes, we examined the genomic evolutionary dynamics of 67 species including at least 5 independent origins of diurnality. Across diurnal spider lineages, hundreds of genes exhibited convergent shifts in selection, including intensified selection on neural, locomotion, and visual system genes, and relaxed selection on several core phototransduction components. Notably, diurnal spiders showed convergent deceleration in evolutionary rates in circadian regulators, such as *CLOCK* and *CRTC1*, and they harbored distinct repertoires of positively selected genes relative to non-diurnal species. In addition, convergent amino acid substitutions were enriched in diurnal hunting spiders. Comparative multi-tissue transcriptomics showed that genes under convergent selection, particularly those involved in vision, sensory processing, nervous system development, and locomotion, tended to exhibit stronger eye- and brain-biased tissue specificity in diurnal species. Altogether, our results reveal convergent genomic changes associated with repeated evolution of diurnality and illustrate how ecological light environments repeatedly shape the molecular evolution of complex animal behavior.

## Introduction

Spiders are among the most abundant and evolutionarily successful arthropod predators, occurring across nearly all terrestrial habitats, from forests and grasslands to deserts, wetlands, caves, intertidal and urban environments ^1,2^. As generalist predators, they have evolved diverse behavioral and physiological traits that allow them to exploit a wide range of prey and environmental conditions ^3,4^. Among these behavioral traits, diel activity (the pattern of activity across the day–night cycle) represents a major axis of behavioral variation and adaptation, that determines when spiders forage, which prey and predators they encounter, and how their behavioral rhythms have evolved in response to ecological and evolutionary pressures. Animals are commonly categorized as diurnal, nocturnal, or crepuscular depending on the hours of day when they are most active ^5^. Although entomologists generally expect that more spiders are active at night and considered nocturnal activity, likely reflecting reduced predation risk and exploitation of nocturnal insect prey ^6–8^. Field observations or laboratory-based assessments reveal remarkable variation in diel activity among species ^6,9^. In several species, males and females differ in their diel activity patterns, reflecting sex-specific ecological or reproductive strategies ^9,10^. Nevertheless, diurnality has evolved repeatedly in multiple spider lineages, such as jumping spiders (family: Salticidae), crab spiders (family: Thomisidae) and lynx spiders (family: Oxyopidae) ^11–13^ (Fig. 1a). These repeated transitions suggest that shifts in diel activity represent adaptations to diverse ecological pressures. Several hypotheses have been proposed to explain the repeated transitions to diurnality, including predator avoidance, prey specialization, and signaling ecology ^5,12,14,15^. However, the mechanism underlying repeated transitions to diurnality remain poorly understood.

**Figure 1.**
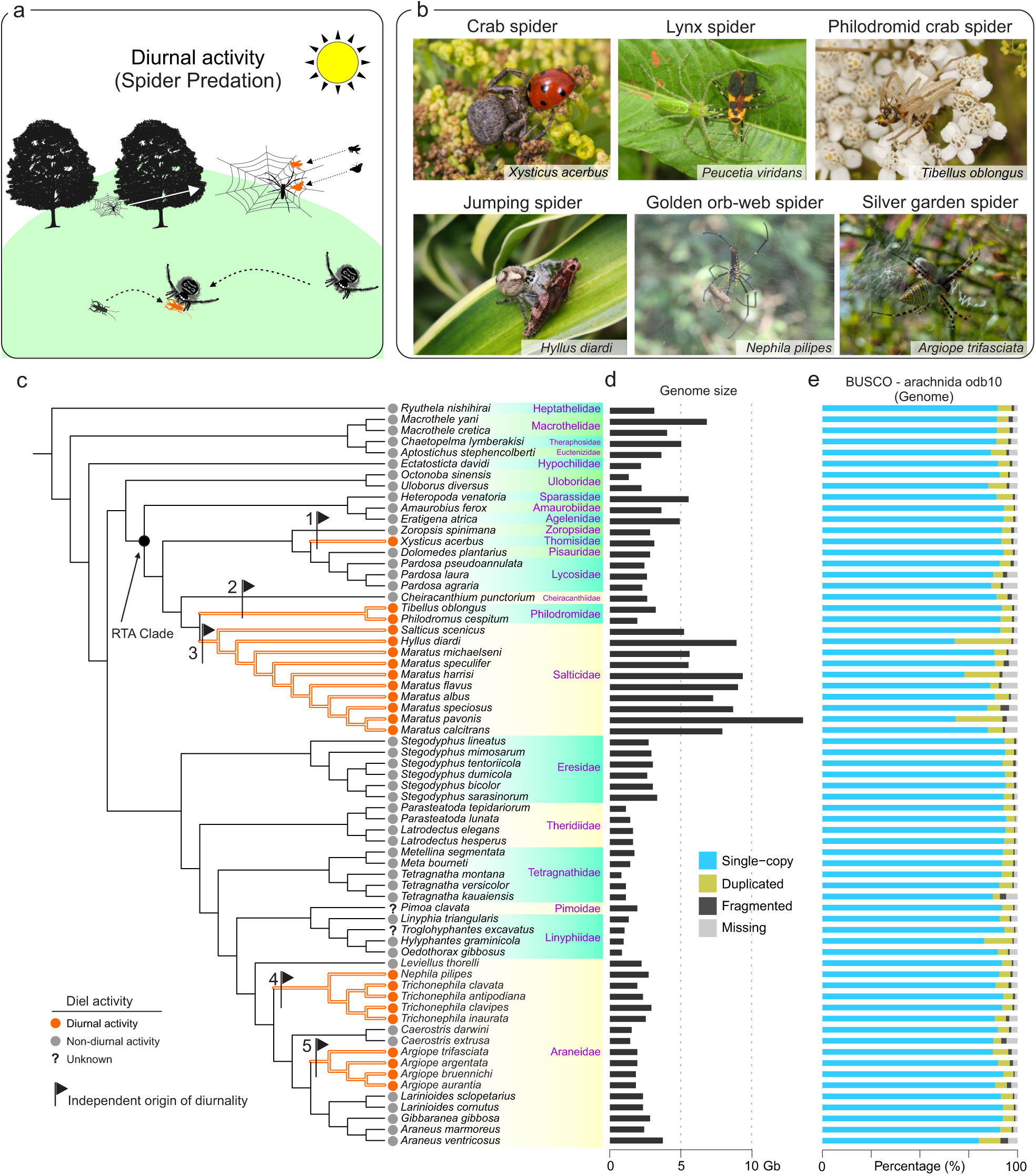
Spider diurnality and genomic features. (a) Schematic diagram depicting the diurnal activity and hunting of spiders. Credit: tree, insects, and spiders are from http://phylopic.org/ (Gabriela Palomo-Munoz [Quercus rubra], Andy Wilson [*Vanessa cardui*], Ferran Sayol [*Megachile mucida*], Guillaume Dera [*Acheta domestica*], Kamil S. Jaron [*Maratus speciosu*], Gabriela Palomo-Munoz [*Trichonephila clavata*]). (b) Pictures of representative diurnal spiders from *Xysticus acerbus* (credit: Raoul Gerend), *Peucetia viridans* (credit: Pedro Alanis), *Tibellus oblongus* (credit: Bob Tong), *Hyllus diardi* (credit: Michał Górski), *Nephila pilipes* (credit: Woodrose Liu), *Argiope trifasciata* (credit: Brooke Washburn). (c) The maximum likelihood (ML) phylogenetic tree of 67 spider species included in the study. The ML tree was inferred from 2,360 core-shared single-copy BUSCO genes. Orange dots represent diurnal spider lineages, grey dots represent non-diurnal spider lineages. Dark flags represent the independent origins of diurnality. (d) Bar plot depicting the genome sizes of 67 spider species. (e) Bar plot depicting the BUSCO profiles of 67 spider genomes, including the percentages of single-copy, duplicated, fragmented, and missing orthologs.

Across animals, diel transitions are widespread and have been extensively studied in vertebrates, where shifts between nocturnality and diurnality typically occur within a highly conserved visual and circadian framework ^16–20^. Unlike vertebrates, spiders possess a multi-eyed and highly specialized visual system, which provides a uniquely complex sensory context for diel transitions ^21^. Previous studies had mainly focused on morphological and behavioral correlates of diel activity patterns in spiders, such as the evolution of enlarged principal eyes in jumping spiders that facilitate visually guided hunting during the day ^21,22^, and the cryptic coloration of crab spiders that enables effective ambush predation on flowers (Fig. 1b) ^23,24^.

Recent large-scale comparative morphological analysis revealed that diurnal spiders tend to have smaller posterior lateral eyes and higher interocular variance, whereas non-diurnal species possess proportionally larger small-eye pairs ^25^. Besides, evidence from other arthropods highlight the role of conserved molecular pathways, including circadian rhythm genes, photoreceptor proteins, and neuromodulatory circuits in regulating diel activity ^26,27^. However, whether similar genetic mechanisms function in spiders is still unclear. Moreover, hunting strategy (web-building versus hunting) represents a key functional trait that influence sensory demands and predator–prey interaction, yet their relationship with diel activity remains unexplored at the genomic level. Thus, it remains unclear whether repeated transitions to diurnality in spiders are caused by shared molecular pathways, or lineage-specific genomic changes acting on sensory and behavioral traits, or both.

Here, we investigated genomic signatures associated with repeated transitions to diurnality in spiders. We combined meta-analysis of published behavioral studies with field observations to assemble a curated dataset of diel phenotypes across the spider phylogeny. Using this behavioral phenotype dataset, we conducted comparative genomics and molecular evolution analyses to identify both convergent and lineage-specific evolutionary changes underlying independent diurnal transitions in spiders. By linking distinct diel activity with gene expression patterns, we finally sought to determine the difference between diurnal and non-diurnal spider at transcriptional level, and especially focused on genes associated with vision, circadian regulation and locomotor activity. Taken together, these analyses provide a comprehensive framework for understanding how ecological context and genomic changes interact to shape complex behavioral traits in one of the most diverse and ecologically significant groups of terrestrial predators.

## Results

### Genome sequencing, assembly, and annotation

To enable robust comparative genomics across independent diurnal lineages, we generated *de novo* assembly for nine diurnal jumping spider (family: Salticidae) genomes using long-read PacBio HiFi sequencing, including newly sequenced HiFi reads for *Hyllus diardi* and publicly available HiFi reads for eight *Maratus* species. To improve genome assembly for *Hyllus diardi*, we generated additional Oxford Nanopore sequencing reads (Fig. 1c; Supplementary Table 1). These newly assembled genomes ranged from 5.51 Gb to 13.59 Gb (Fig. 1d; Supplementary Table 1). Notably, jumping spiders have relatively large genome size (8.094 Gb on average) compared with other spider lineages. BUSCO assessments (arachnida odb10) indicated high assembly completeness, ranging from 90.5% to 96.9% (Fig. 1e, Supplementary Table 1). We annotated 27,000 – 32,000 protein-coding genes per genome, and gene set completeness (BUSCO, arachnida odb10, model: protein) ranged from 90.1% to 94.2% (Supplementary Table 1). These gene counts are comparable to those reported for other spider genomes (Supplementary Table 1).

In addition, we initially incorporated 64 published spider genome assemblies into our comparative dataset, of which 38 genomes had no publicly available gene models (Supplementary Table 1). For these genomes, we performed comprehensive gene prediction using a combination of *ab initio*, homology-based, and transcriptome-based approaches. Together with existing annotations from published spider genomes, BUSCO assessments (arachnida_odb10, model: protein) revealed a relatively high degree of completeness in the predicted gene sets, ranging from 77.3% to 95.5%. Finally, we confirmed 67 spider genomes with high quality BUSCO metrics, detecting at least 90% of arachnid conserved genes (2,934 genes) at both genome assembly (model: genome) and proteome (model: protein) levels, which are robust and suitable for downstream comparative and evolutionary analyses (Supplementary Table 1).

### A genome-scale phylogeny of spiders

To pinpoint diurnal and non-diurnal species within a rigorous evolutionary framework, we reconstructed a genome-scale phylogeny using single-copy orthologs. Specifically, we obtained 2,360 complete BUSCO single-copy orthologs across all 67 spider species and one outgroup species *Centruroides vittatus* (Supplementary Table 1). We reconstructed a phylogenetic tree based on the concatenated single-copy orthologs, which was strongly supported (bootstrap value = 100) for all nodes, and was mostly consistent with previously published spider phylogenies ^28–33^.

### Independent origins of diurnal transition in spiders

To determine how many times diurnality evolved independently, we mapped curated diel activity phenotypic data onto the phylogeny. We compiled observed diel activity information for all 67 spider species through a comprehensive literature survey (Fig. 2a, Supplementary Table 2). Specifically, we initially obtained 457 behavioral studies associated with spider diel activity. After removing duplicates, 314 relevant diel behavioral studies were retained. We finally obtained 109 diel behavioral studies after manual screening with a set of filtrations (Fig. 2a). Among these species, ten from family Salticidae, one from family Thomisidae, two from family Philodromidae, nine from family Araneidae were primarily active during daylight hours and were therefore classified as diurnal spiders (Fig. 2b). Although several spider species exhibited occasional daytime activity, their predominant activity occurred at dusk, dawn, or during the night. These species were thus categorized as nocturnal or crepuscular spiders (hereafter collectively referred to as “nondiurnal” spiders). To further validate the diel activity patterns, we conducted field behavioral observations on four representative spider species in the northeastern United States, including *Tibellus oblongus* (family: Philodromidae), *Salticus scenicus* (family: Salticidae), *Trichonephila clavata* (family: Araneidae), and *Argiope trifasciata* (family: Araneidae) (Fig. 2b, Supplementary Table 3). All four species exhibited active foraging and locomotor behaviors predominantly during the day (Fig. 2b), although *T. clavata* and *A. trifasciata* had occasional activity at other times. Field observations supported our classification of these species as diurnal spiders. By integrating the diel activity dataset with reconstructed genome-scale phylogeny, we identified at least five independent origins of diurnality across 67 species (Fig. 1c). These independent transitions provide a comparative framework for exploring genomic signatures associated with repeated behavioral evolution in spiders.

**Figure 2.**
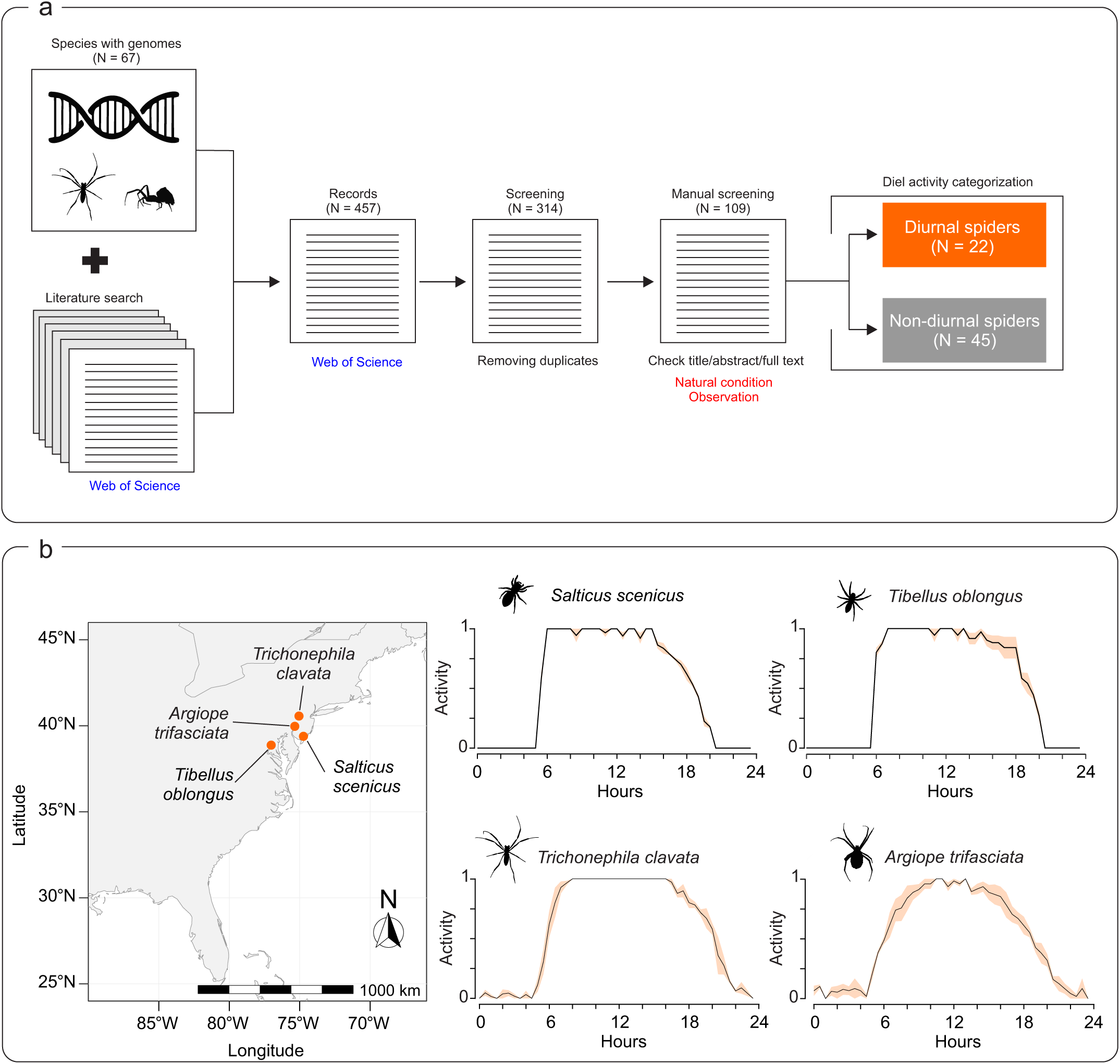
Meta-analysis and field observation of spider diurnal activity. (a) The flowchart depicting the literature search of behavioral studies associated with spider diel activity. (b) Map depicting the geographical locations of four representative diurnal species in northeastern regions of the United States. The time-series activity plot depicting hourly activity levels presents the mean curve with a shaded confidence band (SE as an estimate of the 95% CI). Silhouette illustration of *Trichonephila clavata* from http://phylopic.org/ (credit: Gabriela Palomo-Munoz), *Salticus scenicus*, *Tibellus oblongus* and *Argiope trifasciata* made by Chao Tong.

### Gene-wide relaxation or intensification associated with repeated diunral transition

To test whether shifts in selective pressures are associated with repeated transitions to diurnality, we applied RELAX ^34^ to quantify gene-wide relaxation or intensification of selection. Specifically, we obtained 8,483 one-to-one orthologous genes shared by 67 spider species using FastOMA ^35^ and estimated shifits in selective intensity for each ortholog across the spider phylogeny. We used RELAX ^34^ to identify genes that experienced relaxed or intensified selection in all diurnal species relative to non-diurnal species. We found that more genes experienced intensified selection (453 genes, K > 1, *q*-value < 0.05) than genes under relaxed selection (313 genes, K <1, *q*-value < 0.05). Among genes under relaxation, we identified a set of phototransduction genes, such as r-opsin (K = 0.6641, *q*-value < 0.05) and peropsin (K = 0.6745, *q*-value < 0.05) (Fig. 3a, Supplementary Table 4). In addition, we identified genes involved in movement and locomotion activity under intensification of selection, such as actin-binding LIM protein 2 (ABLIM2) (K = 1.3651, *q*-value < 0.05). Moreover, we found 208 gene ontologies (GO) significantly associated with convergent intensified selection in diurnal spiders were mainly related to neural function, sensory organ and eye development, such as synapse organization (GO:0050808), nervous system development (GO:0007399), visual system development (GO:0150063), sensory organ morphogenesis (GO:0090596), eye development(GO:0001654) (Fig. S1a, Supplementary Table 5). Finally, we found 29 GO terms significantly associated with convergent relaxed selection in diurnal spiders mainly associated with metabolic process and stimulus response, including carbohydrate metabolic process (GO:0005975), biosynthetic process (GO:0009058), and response to stimulus (GO:0050896) (Fig. S1b, Supplementary Table 5).

**Figure 3.**
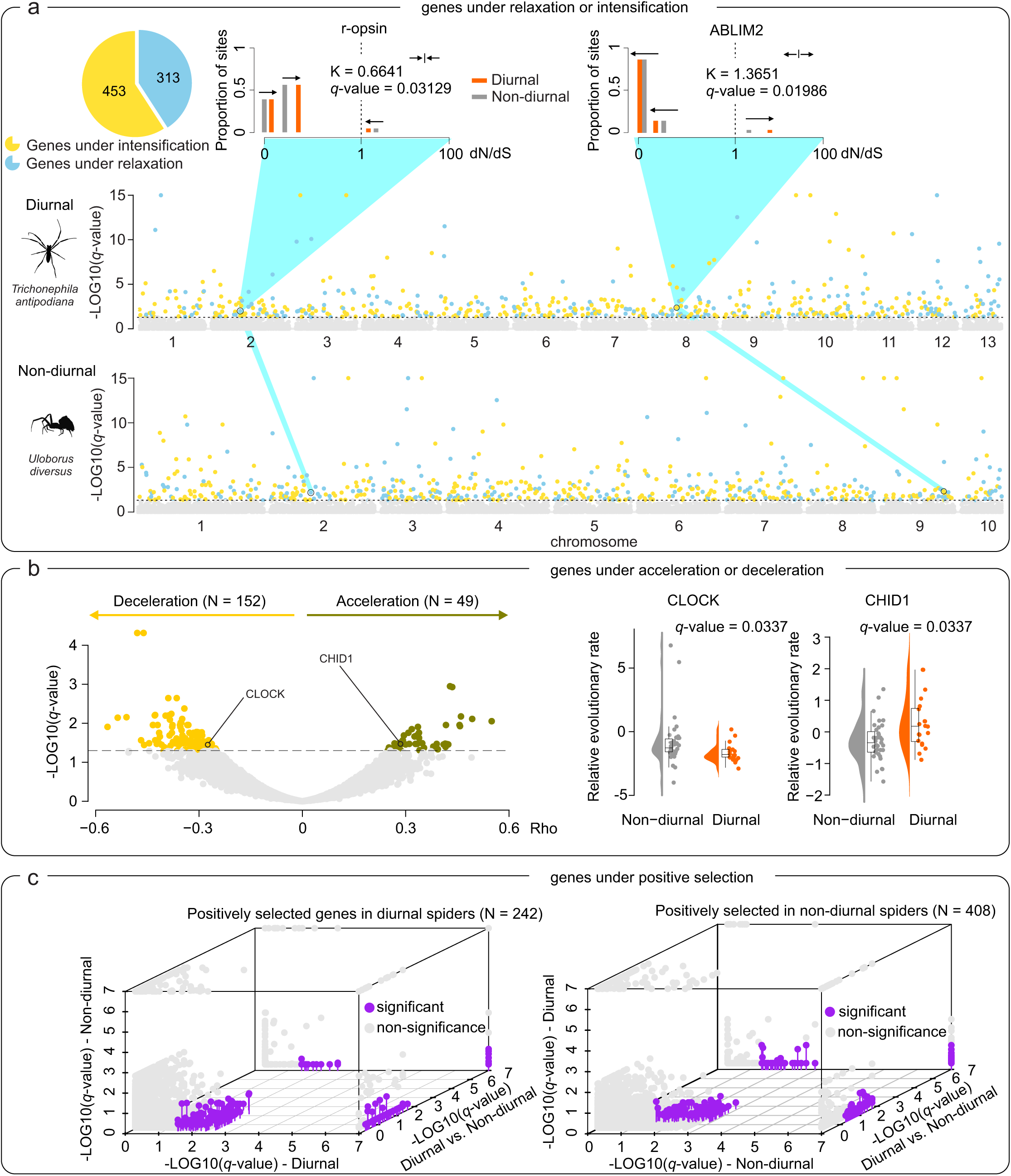
Convergent molecular evolution signatures associated with repeated transitions to diurnality in spiders. (**a**) Pie charts depicting the number of genes under relaxed (bluegreen) or intensified (light yellow) selection associated with repeated transitions to diurnality. Multi-bar plots depicting the patterns of selection experienced across sites in representative gene r-opsin (relaxed selection) and ABLIM2 (intensified selection) of diurnal and non-diurnal spiders, estimated with the Partitioned Descriptive model in RELAX. P value as calculated by using Likelihood-ratio test (LRT). The distribution of dN/dS across sites in the genomes are illustrated by three categories of dN/dS for diurnal (orange) and non-diurnal (grey) branches. The vertical dashed line at dN/dS□=□1 represents neutral evolution, bars at dN/dS□> □1 represent sites experiencing positive selection, and bars at dN/dS <1 represent sites experiencing purifying selection. The arrows show the direction of change in dN/dS between non-diurnal and diurnal branches, demonstrating relaxation of purifying and positive selection associated with the transition to diurnality. K□<□1 indicates relaxed selection, K > 1 indicates intensified selection. Manhattan plots depicting the chromosome distribution of genes under relaxed or intensified selection in representative diurnal spider (*Trichonephila antipodiana*) and non-diurnal spider (*Uloborus diversus*) genomes. The yellow dots represent genes under relaxed selection with significance (*q* value <= 0.05), bluegreen dots represent genes under intensified selection with significance (*q* value <= 0.05), light grey dots represent genes under selection without significance (*q* value > 0.05). Silhouette illustration of *Trichonephila clavata* from http://phylopic.org (credit: Gabriela Palomo-Munoz) and *Uloborus diversus* made by Chao Tong. (**b**) Volcano plot depicting the genes with accelerated (olive) or decelerated (deep yellow) evolutionary rates in diurnal than non-diurnal spider branches with significance (*q* value <= 0.05). Light grey dots represent the genes under selection without significance (*q* value > 0.05). Volin-box plots depicting the relative evolutionary rates (RER) of CLOCK and CHID1 genes in diurnal and non-diurnal spider branches. Each dot represents a species-level RER estimate, and the box plot summarizes the distribution between diurnal and non-diurnal spiders. (**c**) 3D scatter plot depicting the positively selected genes in diurnal spiders (left) and non-diurnal spiders (right). Blue dots depicting the genes under positive selection with significance, and light grey dots represent genes under selection without significance.

### Accelerated or decelerated genes in diurnal lineages

As shifts in selective constraint can manifest as corresponding changes in evolutionary rate ^36^, we next examined whether the rate of molecular evolution differs statistically between diurnal and non-diurnal spiders. We estimated relative evolutionary rate (RER) for each ortholog across all nodes of the phylogeny, and then standardized these rates relative to the genome-wide distribution using RERconverge ^37^. Out of 8,483 one-to-one orthologs included in the analysis, we identified 49 genes showing convergent acceleration in diurnal branches (Rho > 0, *q*-value < 0.05). These genes, which exhibited consistently higher RERs in diurnal than in non-diurnal branches, were defined hereafter as diurnal-accelerated genes. Conversely, we detected 152 genes showing convergent deceleration in diurnal branches (Rho < 0, *q*-value < 0.05), which exhibited consistently lower RERs in diurnal than in non-diurnal branches and were defined as diurnal-decelerated genes (Fig. 3b, Supplementary Table 6). For instance, diurnal-decelerated genes included several circadian genes, such as circadian locomotor output cycles protein kaput (CLOCK, Rho = -0.2796, *q*-value < 0.05, Fig. 3b). Whereas, diurnal-accelerated genes comprised several exoskeleton remodeling and locomotor activity related genes, such as chitinase domain-containing protein 1 (CHID1, Rho = 0.2776, *q*-value < 0.05) and excitatory amino acid transporter 1 (EAAT1, Rho = 0.428715424, *q*-value < 0.05) (Fig. 3b, Supplementary Table 6). A total of 19 GO terms significantly associated with convergent deceleration (*q* value < 0.05) were mainly related to nervous system development and stimulus response functions, including nervous system development (GO:0007399), animal organ development (GO:0048513), and cellular response to stimulus (GO:0051716) (Fig. S2, Supplementary Table 7).

### Distinct positively selected gene repertoires in diurnal and non-diurnal lineages

We further sought to test for association between positive selection and independent diurnal transitions in spiders. By using <u>B</u>ranch-site <u>U</u>nrestricted <u>S</u>tatistical <u>T</u>est for <u>E</u>pisodic <u>D</u>iversification for <u>PH</u>enotype (BUSTED-PH) ^38^, a method to test for evidence of episodic diversifying selection associated with a specific feature/phenotype/trait (i.e., diurnal activity), we detected the signal of positive selection in diurnal spiders and non-diurnal spiders for each ortholog across the phylogeny, respectively. We identified 243 positively selected genes in diurnal spiders, such as CREB-regulated transcription coactivator 1 (CRTC1) associated with circadian rhythm, ataxin 2 (ATXN2) and MYND domain-containing protein 11 (ZMYD11) involved in nervous system development, tubulin-specific chaperone D (TBCD) and actin-binding LIM protein 2 (ABLIM2) involved in locomotion, Arrestin 1 (ARR1) involved in visual system. In addition, we identified 408 positively selected genes in non-diurnal spiders, such as Chaoptin (chp), king tubby (ktub), homeobox protein BarH1 (barH1) involved in visual system, Cyclin-dependent kinase 5 (Cdk5) associated with circadian clock (Fig. 3c, Supplementary Table 8). Intriguingly, we found that diurnal and non-diurnal spiders shared a cluster of significantly enriched GO terms (*q* value < 0.05) with high semantic similarity, such as nervous system development, neuron development, response to stimulus, and metabolic process, even though the two groups of spiders did not share any positively selected genes (Fig. S3, Supplementary Table 9).

### Intersection of genes under convergent selection

To assess whether different evolutionary signals reflect shared genes, we quantified overlaps among genes under relaxation/intensification, acceleration/deceleration, and positive selection (Fig. 4a, Supplementary Table 10). We identified 47 genes that experienced both positive selection and intensified selection (*p* value < 0.01, SuperExact test), representing cases of intensified positive selection, including CRTC1 associated with circadian rhythm, ATXN2 involved in nervous system development, TBCD and ABLIM2 involved in locomotion, ARR1 involved in visual system (Fig. 4b). In addition, 25 genes were found to be under both intensified selection and deceleration in evolutionary rates (*p* value < 0.01, SuperExact test), such as zinc finger MYM-type protein 4 (ZMTM4) associated with nervous system development, integrin alpha-9 (ITGA9) associated with locomotion. We also detected 14 genes that simultaneously experienced relaxed selection and acceleration in evolutionary rates (*p* value < 0.01, SuperExact test), such as chitinase domain-containing protein 1 (CHID1) involved in exoskeleton remodeling (Fig. 4b). Finally, we identified two genes under both accelerated evolution and positive selection, such as ZMYD11 associated with nervous system development (Fig. 4b).

**Figure 4.**
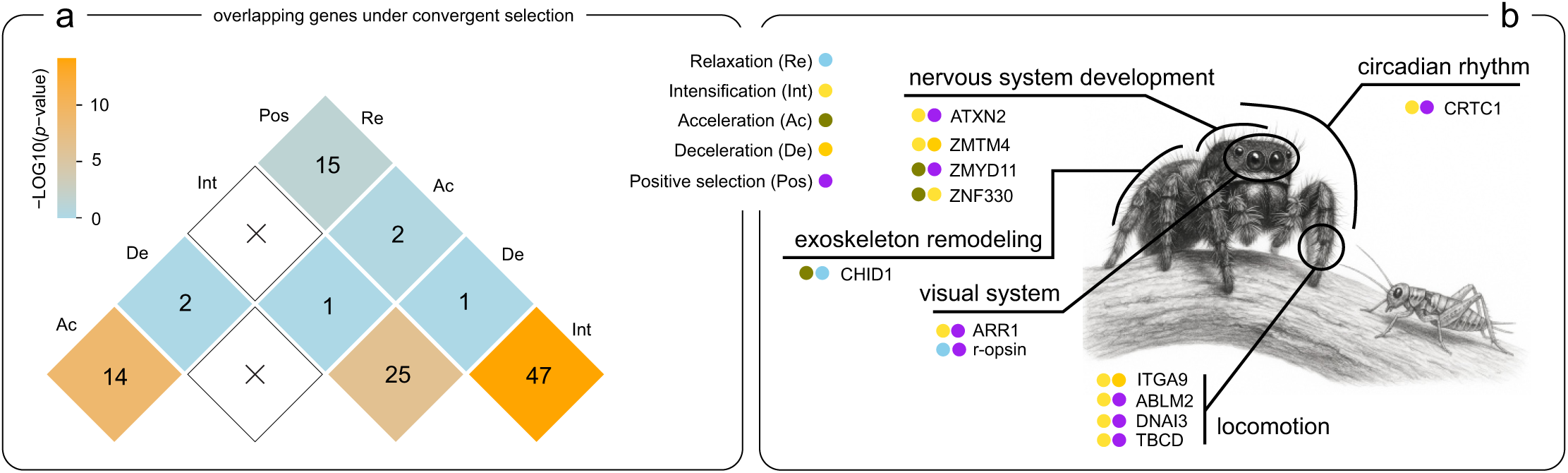
Intersection of genes under convergent selection. (**a**) Pairwise overlap of genes under two distinct selection categories, including relaxation (Re), intensification (Int), acceleration (Ac), deceleration (De), and positive selection (Pos). Each square shows the number of overlapping genes, with color representing the statistical significance of the overlap based on the SuperExact test with log-transformed p value. White square marked with “X” indicates no overlapping genes. (**b**) Overlapping genes involved nervous system development, circadian rhythm, visual system, and exoskeleton remodeling were labeled. The dots in different colors represent the genes under specific selection category. The illustration of a jumping spider and prey was made by Chao Tong.

### Convergent amino acid substitutions associated with repeated diurnal transitions

We extended our analysis to search for convergence in amino acid substitution site (CAAS) associated with repeated transitions to diurnality. We used CAAStools ^39^ to identify convergent amino acid substitutions (CAAS) in diurnal spiders compared to non-diurnal spiders. After filtering candidate sites according to the CAAS criteria, we did not identify any positions where all diurnal species shared one identical amino acid and all non-diurnal species shared another (Pattern 1 sites in CAAStools). Nevertheless, we detected a substantial number of sites at which all diurnal spiders shared an identical amino acid, whereas non-diurnal spiders exhibited diverse residues at the same positions (Pattern 2 sites in CAASTools) (Fig. S4 and Supplementary Table 11).

We further performed lineage-specific CAAS analyses in two subgroups: web-building spiders and hunting spiders. In hunting spiders, we identified 27 “Pattern 1 sites” in 27 genes, as classified by CAAStools, such as Vacuolar protein sorting-associated protein 35 (VPS35) ^40^. In contrast, only a single “Pattern 1 site” was detected in Peroxisome biogenesis factor 10 (PEX10) among web-building spiders (Fig. S4 and Supplementary Table 11).

### Distinct gene expression patterns between diurnal and non-diurnal spiders

We finally sought to quantify the degree of divergence in gene expression patterns between diurnal and non-diurnal spiders. We compared the expression patterns of genes under selection in six tissues of representative diurnal spider (*Hyllus diardi*) and non-diurnal spider (*Heteropoda venatoria*), including posterior lateral eye (PLE), posterior median eye (PME), anterior lateral eye (ALE), anterior median eye (AME), brain, and leg muscle (Fig. 5a).

**Figure 5.**
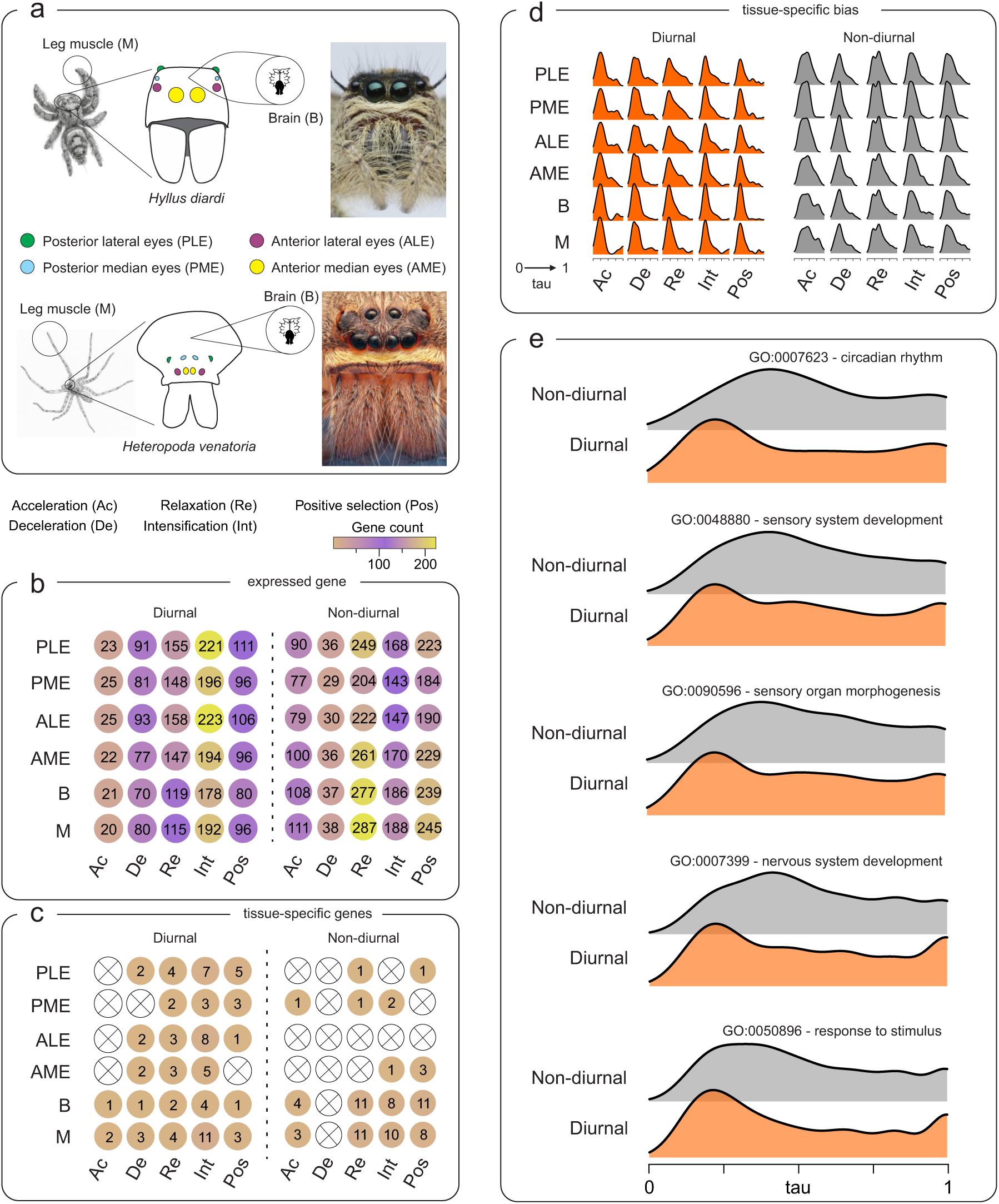
Distinct gene expression patterns between diurnal and non-diurnal spiders. (**a**) Experiment design of comparative transcriptomic analysis. Four eye pairs, brain and leg muscle were sampled from representative diurnal spider *Hyllus diardi* and non-diurnal spider *Heteropoda venatoria*. (**b**) Multi-dot plots depicting the number of genes under different selection categories that expressed in each spider tissue. (c) Multi-dot plots depicting the number of genes under different selection categories that are specifically expressed in a unique tissue. (**d**) Ridgeline plots depicting the tissue-specific bias of under different selection categories based on tau value. (**e**) Ridgeline plots depicting the tissue-specific bias of genes involved in circadian rhythm, sensory system development, sensory organ morphogenesis, nervous system development, response to stimulus in diurnal and non-diurnal spiders.

Across five selection categories, two spiders exhibited distinct gene expression pattern at tissue-level (Fig. 5b, Supplementary Table 12). For accelerated genes, diurnal spiders showed largely homogeneous expression across tissues, whereas non-diurnal spiders expressed fewer genes in PME and ALE. For decelerated genes, diurnal spiders showed enrichment in PLE and ALE, while non-diurnal spiders exhibited no detectable tissue differences. For genes experiencing relaxed selection, diurnal spiders expressed substantially more genes in four eye tissues than in brain or leg muscle, whereas PME and ALE harbored fewer expressed genes in non-diurnal spiders. A similar pattern was observed for genes under intensified selection: PLE and ALE in diurnal spiders harbored more expressed genes, whereas PME and ALE in non-diurnal spiders included fewer. For positively selected genes, diurnal spiders expressed the most in PLE and ALE, in contrast to non-diurnal spiders, which expressed fewer in PME and ALE and the most in the leg muscle.

We next identified tissue-specific genes, defined as those expressed exclusively in a single tissue (Fig. 5c, Supplementary Table 13). Accelerated genes showed little tissue specificity in either spider eye tissues, except for one PME-specific gene in non-diurnal spiders. Diurnal spiders harbored multiple tissue-specific genes for deceleration, relaxation, and intensification categories, whereas non-diurnal spiders showed reduced tissue specificity, lacking such genes in key eye tissues (e.g., ALE, AME, PLE) under several selection types. For positively selected genes, diurnal spiders lacked AME-specific genes, while non-diurnal spiders lacked PME- and ALE-specific genes.

Further, we assessed how selection influences expression specialization and analyzed the tissue-specific bias of genes across six tissues. Tissue-specific bias describes the extent to which a gene is preferentially or exclusively expressed in one tissue, measured using tau (τ), where values close to 1 denote strong tissue specificity and values near 0 indicate broad expression. Although the tau distributions varied across tissues within each selection category, the overall expression pattern revealed a clear difference between the two groups: in diurnal spiders, genes under selection tended to be more exclusively expressed, whereas in non-diurnal spiders these genes were generally more broadly expressed (Fig. 5d, Supplementary Table 14).

Finally, we focused on genes associated with several enriched functional categories that repeatedly highlighted in previous GO enrichment outcomes, including circadian rhythm (GO:0007623), sensory system development (GO:0048880), nervous system development (GO:0007399), and response to stimulus (GO:0050896). Genes associated with these functional categories showed distinct patterns between two spider groups. In diurnal spiders, they tended to be expressed in a more tissue-specific manner, whereas in non-diurnal spiders they were more broadly expressed across tissues (Fig. 5e, Supplementary Table 15).

## Discussion

The repeated transition to diurnality from predominantly nocturnal or crepuscular ancestors represents a striking example of behavioral convergence in spiders. By integrating the largest comparative genomic dataset for spiders to date, curated diel activity phenotypes derived from literature synthesis and field observations, genome-wide test of molecular evolution, and tissue-specific transcriptomic profiling, we revealed convergent and lineage-specific genetic changes associated with at least five independent transitions to diurnal activity.

Transitions from nocturnal or crepuscular activity to a diurnal lifestyle represent dramatic behavioral shifts that substantially reshape a spider’s visual ecology, altering both prey availability and predation risk. Such ecological changes imply that the visual systems of diurnal spiders may experience strong selective pressures to accommodate brighter, more variable daytime environments ^21,41^. By comparing the genome-wide evolutionary dynamics of multiple independently evolved diurnal lineages with those of their closely related non-diurnal species, we identified a set of phototransduction, and visual-system associated genes that experienced natural selection. For example, the gene *r-opsin*, which experienced intensified positive selection in diurnal spider lineages, regulates Gq-mediated phototransduction that enables light detection and visual perception in arthropods ^42,43^ and knock-down *r-opsin* in moths exhibited reduced phototactic efficiency to green light ^44^. The gene *Arrestin 1* (*Arr1*), which also experienced intensified positive selection in diurnal spider lineages, encodes a protein involved in rhodopsin inactivation that contributes to photoreceptor maintenance ^45^. The knock-down of *Arr1* in fly can alter the termination of rhodopsin signaling and Rh1 endocytosis, enhancing light responses and predisposing photoreceptors to light-dependent degeneration ^46^. These genomic signatures are putatively associated with several known behavioral difference between diurnal and non-diurnal spiders. For instance, diurnal spiders (i.e. jumping spider, family: Salticidae) exhibit high-acuity, color-sensitive vision and rely heavily on complex visual cues for prey detection, mating displays, and navigation ^21,47,48^. In contrast, non-diurnal spiders (i.e. giant crab spiders, family: Sparassidae) depend more on mechanosensation and low-light motion detection ^49,50^. In addition, the biased tissue specificity of vision-related genes in diurnal spider is therefore consistent with their highly specialized anterior median eyes (AME) and lateral eyes (PLE, ALE), which support color discrimination, high temporal resolution, and motion tracking during daylight ^21,51^.

Shifting activity into daylight exposes spiders to strikingly different visual environments, prey communities, thermal conditions, and predation pressures ^15,52,53^. These changes require not only behavioral flexibility but also coordinated modifications to endogenous circadian rhythm that synchronizes sensory, neural, and metabolic processes to external diel cycles, the differences between diurnal and non-diurnal spiders had long been documented by past behavioral and physiological studies ^54,55^. Notably, our comparative genomics analysis reveal that this behavioral transition is accompanied by repeated evolutionary changes in circadian components. The core circadian pacemaker gene *CLOCK* exhibits significant convergent deceleration in evolutionary rate in diurnal spiders, indicating relaxed purifying selection on the central transcriptional activator that drives the period–timeless negative-feedback loop in arthropods ^56,57^. The deceleration is consistent with reduced functional constraint on strict 24-hour periodicity when activity occurs under the prolonged and predictable photophase of daylight, where precise endogenous timing putatively becomes less critical for synchronizing physiology and behavior. In addition, the downstream circadian regulator *CRTC1* shows strong signatures of intensified positive selection exclusive to diurnal lineages. *CRTC1* modulates the amplitude of clock-controlled behavioral outputs and integrates environmental light and metabolic cues onto CREB-dependent transcription in fly ^58^. Convergent selection on *CRTC1* is therefore interpreted as adaptive tuning of circadian pathways that support continuous arousal and foraging under the extended daylight conditions experienced by diurnal spiders. Thus, the combination of convergent deceleration of core circadian oscillator and intensification of its downstream regulation pathways, diurnal spiders may repeatedly recruit an ancestral non-diurnal clock into a flexible, light-responsive system that relieves behavior from strict periodicity and optimize sustained activity under the predictably illuminated conditions of daytime predation.

In addition to repeated evolution of spider visual systems and circadian rhythm, behavioral transition to diurnality consistently imposed intense demands on rapid sensory integration, precise motor coordination, and real-time decision-making in complex habitats. Such ecological pressures imply that nervous systems of diurnal spiders may experience strong convergent selection to support the high-speed processing and motor precision required for active hunting in daytime ^11,21,59^. Consistent with these elevated neural and locomotor requirements, our genome-wide evolutionary analyses detected convergent signatures of intensified positive selection on genes regulating nervous system development, synaptic transmission, and cytoskeletal dynamics. For example, the gene *ABLIM2* which experienced strong positive selection in diurnal lineages, plays an essential role in axon guidance and dendritic arborization in developing neurons ^60^, whereas gene *TBCD* is critical for microtubule polymerization and stability during neural circuit formation ^61^. Similarly, the genes *ATXN2* and *ZMYND11*, both under convergent positive selection, regulate RNA metabolism and chromatin organization to modulate neuronal excitability and synaptic plasticity in Drosophila and vertebrates ^62,63^. These genomic signatures directly correspond to well-documented behavioral differences between diurnal and non-diurnal spiders. For instance, diurnal hunters such as jumping spiders (family: Salticidae) and lynx spiders (family: Oxyopidae) execute visually guided pursuits, precise three-dimensional leaps, and mid-air trajectory corrections, exhibit behaviors that are absent in most nocturnal web-building and hunting spiders that rely primarily on vibration or chemosensory cues ^11,21,59^. Moreover, pronounced tissue-specific expression of these neural- and locomotion-related genes in brain and secondary eyes of diurnal *Hyllus diardi* is consistent with enhanced integration of visual input and motor output in specialized vision-motor centers. Thus, convergent intensification of neural circuitry and cytoskeletal effectors emerge as a third major axis, complementary to visual specialization and adaptive circadian rhythm, underlying the repeated evolution of a high-performance, visually driven predatory lifestyle in spiders.

Although our phenotypic classification integrated extensive literature curation and field validation of representative species, some orb-weaving Araneidae included as diurnal exhibit crepuscular tendencies in certain populations, potentially introducing modest noise into convergence tests. Similarly, transcriptomic comparisons are presently limited to one diurnal (*Hyllus diardi*) and one nocturnal (*Heteropoda venatoria*) species, raising the possibility that some expression differences reflect lineage-specific rather than diel-activity-related effects. Future work integrating additional independent transitions to diurnality, multiple developmental stages, single-cell atlases of eye and brain tissues, and behavioral experiments under controlled light environments will further clarify how genomic changes are converted into neural and physiological functions during the evolution of diurnal activity.

In summary, our results provide evidence for coordinated genomic changes associated with repeated phenotypic shifts toward diurnality. These independent transitions consistently involved shared modifications to visual systems, circadian regulation, sensory processing, and neural–motor systems. More broadly, this study illuminates the genomic basis of complex behavior evolution and demonstrates how environmental light conditions can repeatedly shape the molecular mechanisms underlying behavioral adaptation.

### Methods Study design

Spiders exhibit diverse diel activity patterns. Although most species are predominantly nocturnal, diurnality has evolved repeatedly across distantly related lineages. Here, we sought to identify the genomic basis underlying the repeated transitions to diurnality in spiders. We aimed to analyze and compare the genomes of species representing multiple independent origins of diurnality across the spider phylogeny.

### Diel activity date collection

To clarify the diel activity patterns of spider species included in this study under natural conditions, a comprehensive search of the scientific literature was performed using the Web of Science database up until 30 December 2024. The keywords were ‘(spider) AND (diel activity OR nocturnal OR diurnal OR diel rhythm) AND (day OR night) AND (foraging OR locomotion activity OR web-building OR hunting)’. The search returned 457 studies. Titles, abstracts, and keywords were first screened for relevance, and full texts of potentially relevant publications were manually reviewed. The selection process followed PRISMA guidelines and is illustrated in a flow diagram (Fig. 2a). Specifically, we included studies if they (1) examined or observed diel activity in spiders under natural or semi-natural conditions, (2) explicitly reported or quantified activity across both day and night periods, and (3) provided data that could be extracted (e.g., counts of active individuals, predation frequency, or standardized activity indices). Studies conducted solely under laboratory light/dark cycles, without natural diel variation, were excluded, as were those that did not clearly separate day and night observations.

### Study site and field behavioral observation

We selected four spider species which distributed in the northeastern United States, each representing an independent diurnal lineage. Species occurrences were first confirmed by referencing distribution records from the Global Biodiversity Information Facility (GBIF, https://www.gbif.org/). Based on these records, we chose four field sites in July, 2025 to monitor diel activity, including *Tibellus oblongus* (East Potomac Park, Washington, DC, USA), *Salticus scenicus* (Estell Manor Park, Hamilton, NJ, USA), *Trichonephila clavata* (Peace Valley Park, Doylestown, PA, USA), and *Argiope trifasciata* (Veterans Memorial Park, Broomall, PA, USA). At each site, we conducted continuous behavioral observations of foraging and locomotor activity from 08:00 until 08:00 the following day, with observations recorded every 30 minutes.

Each observation session was repeated over three consecutive days, all under clear weather conditions. During each 30-minute interval, we categorized individuals as active or inactive based on their observed behaviors. Active behaviors included walking, jumping, approaching prey, head turning or body movement (environmental scanning), exploration, courtship displays, and stalking or pouncing. In contrast, individuals were classified as inactive when they remained motionless, stayed at the same position for ≥ 5 s, or adopted a resting posture with the body pressed against the substrate and legs extended. For each observation period, we recorded the number of active individuals and calculated the proportion of active individuals (active / total observed) as the activity index. These indices were used to construct diel activity curves for each species.

### Sample collection and genome sequencing

We collected 10 female individuals of a jumping spider species, *Hyllus diardi* from Xishuangbanna, Yunnan, China in June, 2024. Genomic DNA was extracted from a randomly selected female individual using Qiagen Genomic DNA Kit (Cat#13323, Qiagen). The quality and quantity of the extracted gDNA were evaluated using a NanoDrop One UV-Vis spectrophotometer (Thermo Fisher Scientific) and a Qubit 3.0 Fluorometer (Invitrogen), following the manufacturers’ protocols. We prepared a PacBio HiFi library and a Oxford Nanopore library, which were sequenced on a PacBio Sequel II platform and a PromethION platform, respectively.

### Tissue dissection and RNA sequencing

We collected 10 female individuals of a giant crab spider species, *Heteropoda venatoria* from Shenzhen, Guangdong, China in June, 2024, representing a non-diurnal spider lineage. Together with previously collected individuals of the jumping spider *Hyllus diardi*, representing a diurnal lineage, we randomly selected one individual from each species for dissection of 6 tissues, including posterior lateral eyes, posterior median eyes, anterior lateral eyes, anterior median eyes, brain, and muscle from legs. Total RNA was extracted from each tissue using TRIzol (Thermo Fisher Scientific, USA). We assessed the RNA quality with Agilent Bioanalyzer 2100 (Agilent Technologies, USA) using an RNA 6000 Pico Kit (Fisher Scientific, USA), and detected RNA quality with Nanodrop 1000 (NanoDrop Technologies, USA). Library were constructed using the NEBNext Ultra RNA Library Prep Kit and sequenced on a single lane of an illumina NovaSeq 6000 platform.

### Genome assembly

The adapter-contaminated and low-quality Nanopore reads were detected and filtered using NanoPlot ^64^, Porechop_ABI ^65^, and NanoFilt ^66^. We used HiFiAdapterFilt v2.0 ^67^ to perform quality control for PacBio HiFi reads. We performed *de novo* genome assembly for *Hyllus diardi* with both Oxford Nanopore reads and PacBio HiFi reads using Verkko ^68^ with default parameters. In addition, we collected PacBio HiFi reads for other 8 jumping spider species from NCBI SRA (https://www.ncbi.nlm.nih.gov/sra), including *Maratus michaelseni*, *Maratus speculifer*, *Maratus harrisi*, *Maratus flavus*, *Maratus albus*, *Maratus speciosus*, *Maratus pavonis*, *Maratus calcitrans* (Supplementary table1). The adapter-contaminated PacBio HiFi reads were detected and filtered using HiFiAdapterFilt v2.0 ^67^, the remaining PacBio HiFi reads were assembled into contigs using Hifiasm 0.16.0 ^69^. Potential haplotype redundancies in the genome drafts were resolved using Purge_dups v1.0.1 ^70^ with default settings. Finally, we performed the quality of completeness assessment of genome assemblies using BUSCO v5.4.5 ^71^ with arachnida_odb10 data set (N = 2,934).

### Genomic data collection

We collected publicly available spider genome assemblies from online databases, including NCBI Genome (https://www.ncbi.nlm.nih.gov/genome), CNGBdb (https://db.cngb.org/), and GigaDB (http://gigadb.org/), as of December 2024. To ensure data quality and consistency, we recorded assembly statistics such as genome size and BUSCO completeness, when available. All published genomes were classified into two categories: annotated (with publicly available gene models) and unannotated (without gene annotations). For the genomes with no publicly available gene models, we performed gene annotation following the standardized pipeline described below.

### Genome annotation

We identified repetitive elements in each genome using a combination of *ab initio* and homology-based approaches. First, we constructed the species-specific repeat database (*ab initio* database) using RepeatModeler v2.0.1 ^72^. Second, we merged the known repeat library (Repbase, https://www.girinst.org/repbase/) with *ab initio* database as the reference repeat library. Finally, repetitive elements were annotated in each genome using RepeatMasker v4.2.1 ^73^ based on this combined reference.

We annotated gene models following the pipeline described in our previous study ^74^, integrating *ab initio*, homology-based, and transcriptome-based prediction approaches using MAKER v3.01.04 ^75^. For *ab initio* prediction, gene structures were generated using AUGUSTUS v3.3.3 ^76^ and GeneMark-ES/ET/EP v4.48_3.60 ^77^, both of which were pre-trained with BRAKER v3.0.2 ^78^ to improve prediction accuracy. For homology-based prediction, protein sequences from seven representative arthropod and chelicerate species, including *Stegodyphus mimosarum* (GCA_000611955.2), *Parasteatoda tepidariorum* (GCA_000365465.3), *Trichonephila clavipes* (GCA_002102615.1), *Drosophila melanogaster* (GCA_000001215.4), *Ixodes scapularis* (GCA_002892825.2), *Daphnia pulex* (GCA_900092285.2), and *Strigamia maritima* (GCA_000239455.1), which were retrieved from NCBI Genome database (https://www.ncbi.nlm.nih.gov/home/genomes/) and provided to the MAKER pipeline as protein homology evidence. When transcriptome data were available or sequenced, RNA-seq reads were aligned to the corresponding genome using HISAT2 v2.2.0 ^79^ and and assembled into transcripts with StringTie v2.1.3 ^80^. The assembled transcripts were supplied to the MAKER pipeline via the *est* option to refine gene structure annotation.

To evaluate the quality of annotated genomes, we employed the BUSCO pipeline to assess the completeness of annotated proteomes based on arachnida_odb10 data set (N = 2,934). Only genomes with annotated proteomes achieving > 90% complete single-copy BUSCOs were retained for comparative analyses.

### Genome-scale phylogeny construction

We downloaded the genome of striped bark scorpion, *Centruroides vittatus* (GCF_030686945.1) for phylogeny construction as the requirement of an outgroup species. Building on the identified BUSCO gene repertoire (arachnida_odb10) within two categories “Complete and single-copy BUSCO” and “Complete and duplicated BUSCO” of 67 spider genomes and an outgroup genome, we identified one-to-one single copy genes across all 16 species using OrthoFinder ^81^. Next, we used a phylogenomic approach to reconstruct the phylogeny of 67 spider species, along with an outgroup species, based on a dataset of amino acid (AA) sequences corresponding to a pooled set of 1:1 single-copy orthologs. Further, we performed AA sequence alignment using MAFFT ^82^, removed gaps using trimAL ^83^, and assembled a concatenated dataset that included all 1:1 single-copy orthologs with a minimum length of 200 AA. Finally, we used ModelFinder ^84^ to determine best-fit model of sequence evolution and constructed the maximum likelihood (ML) phylogenetic tree using IQ-TREE2 ^85^ with 1000 bootstrap replicates.

### Ortholog identification

BUSCO gene repertoire which only included curated single-copy orthologs from OrthoDB database, we employed FastOMA ^35^ to extend and explore orthologous relationship of protein-coding genes across 67 spider species. Briefly, we compiled all proteome dataset for each spider species and the newly reconstructed genome-scale phylogeny as the requirements.

### Analysis of codon substitution rate associated with diurnality

To quantify the degree to which shifts in the selection during repeated transition to diurnality in spiders, we sought to detect the changes in nucleotide substitution rate (ω) and the selective strength (relaxation or intensification) acting on repeated diurnal transition. Specifically, we prepared the codon alignments of shared orthologs by 67 spider species, which derived from amino acid alignments and corresponding DNA sequences using PAL2NAL v.14 (-no gap) ^86^. We retained the codon alignments with a minimum length of 50 codons, and prepared the corresponding tree for each ortholog by pruning the genome-scale phylogeny using R package, phytools ^87^. Finally, we used RELAX ^34^ to test for relaxation or intensification of selective pressure along diurnal spider branches across the phylogeny. Briefly, RELAX distinguishes between the signals by modeling how codons with different ω categories (ω[>[1 and ω[<[1) respond to a single selection intensity parameter K. Relaxation of selection would push all ω categories toward 1, while intensification of selection would pull all ω categories away from 1. K > 1 indicates the signature of intensified selection, whereas K < 1 indicates a relaxed selection strength at diurnal spider branches. We employed a log-likelihood ratio test (LRT) to compare the supports for the null model (K = 1) and the alternative model (K > 1 or K < 1), which further corrected p value using the Benjamini-Hochberg method to control for multiple comparisons. We defined the genes under convergent relaxation which showed K < and *q* < 0.05, while genes with K >1 and *q* < 0.05 are considered to be under convergent intensification during transition to diurnality.

### Analysis of relative evolutionary rate associated with diurnality

To test whether signature of molecular evolution differs between non-diurnal and diurnal spiders, we computed the relative evolutionary rate (RER) for each species at gene-wide scale. Specifically, we aligned the AA sequences of shared orthologs for the 67 spider species using MAFFT ^82^. Next, we inferred gene tree and estimate branch length for each ortholog by pruning the reconstructed genome-wide phylogeny using R package, Phangorn ^88^ with the AA alignments. We calculated the relative RER for each node on gene tree using R package, RERconverge ^37^, and compared the RERs between non-diurnal and diurnal spiders with the Wilcoxon rank sum test. In addition, we corrected the reported *p* values for multiple comparisons by computing *q* values. Finally, we defined the genes with RERs for diurnal spiders significantly higher than RERs representing non-diurnal spiders as rapid evolving genes or diurnal-accelerated genes (*q* < 0.05).

### Analysis of positive selection associated with diurnality

To determine whether convergent positive selection associated with the repeated transition to diurnality, we sought to detect the signal of positive selection in diurnal spider branches across the phylogeny using BUSTED-PH ^38^ with the codon alignments and gene trees for each shared ortholog as described above. We used LRT to calculate *p* values for focal foreground branches (i.e. diurnal spiders), background branches (i.e. non-diurnal spiders), and difference between focal foreground and background following Benjamini–Hochberg adjustment. We defined genes with significant signals of convergent positive selection in diurnal spider branches (*q*_diurnal_ < 0.05) and difference between diurnal and non-diurnal branches (*q*_diff_ < 0.05), while no significant signals in non-diurnal branches (*q*_non-diurnal_ < 0.05) as positively selected genes in diurnal spiders.

### Analysis of convergent amino acid substitutions associated with diurnality

To test whether specific gene harbor convergent amino acid substitutions (CAAS) in diurnal species relative to non-diurnal species across the phylogeny, we sought to identify CAAS using CAAStools ^39^ with shared ortholog dataset. We first conducted a genome-wide CAAS search by comparing all 22 diurnal species against 45 non-diurnal species. To further evaluate putatively lineage-specific convergence, we divided the spiders into two subgroups based on their hunting models: web-building spiders (45 species) and hunting spiders (22 species). For each subgroup, we performed independent CAAS searches, comparing 9 diurnal versus 36 non-diurnal species within the web-building spider group, and 13 diurnal versus 9 non-diurnal species within the hunting spider group. We applied a strict criterion that defined convergence only when all diurnal species shared an identical amino acid (e.g. D) at a specific site, whereas all non-diurnal species carried another identical amino acids (e.g. E) at the same position.

### Analysis of gene expression patterns between diurnal and non-diurnal spiders

To determine whether diurnal and non-diurnal spiders differ in their gene expression patterns, we conducted a muti-tissue transcriptomic analysis. We first performed quality control on newly sequenced RNA-seq data from six tissues of diurnal spider *Hyllus diardi* and nocturnal spider *Heteropoda venatoria* using fastp ^89^. Next, we mapped these clean reads to spider genome using kallisto (https://pachterlab.github.io/kallisto/) to obtain transcript abundance estimates in Transcripts Per Million (TPM). Furthermore, we assessed gene expression specificity across tissues by calculating τ (tau), a widely used metric of expression specificity ^90^. Tau ranges from 0 to 1, which 0 indicates that genes are ubiquitously expressed and 1 indicates that genes are exclusively expressed in one tissue.

### Gene ontology enrichment analysis

We performed gene ontology (GO) enrichment analysis for these genes under convergent relaxation, convergent acceleration/deceleration, convergent positive selection using GOTermFinder ^91^ and further corrected the reported *p* values following Benjamini–Hochberg adjustment ^92^. We considered the GO terms (Biological Processes) with *q* value < 0.05 to be shown significant enrichment.

## Competing interests

The authors declare no competing interests.

## Author contributions

C.T. and Z.S.Z designed research; C.T., Z.F., and L.Y.W. performed research; C.T. analyzed data; and C.T. wrote the paper.

## Data availability

Raw and processed genome and transcriptome data have been deposited in NCBI SRA (accession number: PRJNA1338085).

## Code availability

All scripts required to perform all analyses are available at Github (https://github.com/jiyideanjiao/spider_diel_activity).

## Acknowledgements

This work was funded by the National Natural Science Foundation of China (3257039), the Special Investigation and Classification of Invertebrates from Yintiaoling Nature Reserve (CQS24C00333), the Science Foundation of School of Life Sciences SWU (20232008071901 and 20212020110501) to Z.S.Z and L.Y.W.

## Notes

### Competing Interest Statement

The authors have declared no competing interest.

